# The synaptic vesicle priming protein Munc13 mediates evoked somatodendritic dopamine release

**DOI:** 10.64898/2025.12.23.696181

**Authors:** Joseph J. Lebowitz, Aditi Banerjee, Gillian Handy, John T. Williams, Pascal S. Kaeser

## Abstract

Midbrain dopamine neurons secrete dopamine from their somata and dendrites in addition to their axonal release. Shared and distinct properties have been proposed for somatodendritic and axonal release, but the mechanisms of somatodendritic release remain unclear. We here used mouse genetics, electrophysiology, and imaging to define roles of the synaptic vesicle priming protein Munc13 in somatodendritic release in comparison to axonal secretion. Munc13 ablation in dopamine neurons decreased evoked but not spontaneous somatodendritic dopamine transmission measured as D2 receptor-mediated currents. Imaging with a fluorescent dopamine sensor confirmed this finding and established comparable importance for Munc13 in somatodendritic and axonal secretion. Pharmacological experiments revealed that release from adrenergic terminals contributes to D2 receptor-mediated currents, and their relative contribution was enhanced after Munc13 knockout. Altogether, these data establish important roles for Munc13 in evoked somatodendritic release. These roles are similar to Munc13 functions in axonal dopamine release and at fast synapses. Spontaneous midbrain dopamine release does not necessitate Munc13 in dopamine neurons and may rely on a release pathway that is independent of the prototypical release machinery employed at synapses.

## Introduction

Neurotransmitter release from axon terminals is mediated by specialized secretory machinery for ultrafast vesicle fusion ^1,2^. Most neurons also release neurotransmitters from the somatodendritic compartment, and both synaptic point-to-point transmission and volume transmission that lacks synaptic structure may mediate this dendro-dendritic signaling ^3^. Midbrain dopamine neurons, for example, release dopamine from somata and dendrites, and D2 receptor activation inhibits their firing ^4–8^.

Recent work has started to identify molecular machinery for somatodendritic dopamine release. First studies established that release is vesicular and relies on SNARE proteins for exocytosis ^9–11^. Evoked somatodendritic release is Ca^2+^-dependent and utilizes Synaptotagmin-1 as a Ca^2+^ sensor ^10,12^, and additional Ca^2+^ sensors might regulate it ^13–16^. Voltage-gated Ca^2+^ channels are a Ca^2+^ source, though the precise identity of the channels remains uncertain ^10,16^. Furthermore, the active-zone protein RIM is important for stimulus-induced somatodendritic dopamine release, suggesting that preorganized protein complexes tethered by RIM mediate these fusion events similar to axonal dopamine secretion ^17,18^. In aggregate, these studies revealed that the molecular machinery for somatodendritic dopamine release resembles that of classical synaptic release, supporting the model that small, clear vesicles might be the fusing membrane compartment ^19,20^.

Spontaneous somatodendritic dopamine release occurs in the absence of action potentials, is Ca^2+^-independent, and does not rely on RIM or Synaptotagmin-1 in dopamine neurons ^12,18,21^. While readily detected in somatodendritic regions as miniature events recorded as D2 receptor-mediated inhibitory postsynaptic currents (D2-IPSCs), it has remained uncertain whether this form of release is also present in axons. Ablating RIM from dopamine neurons does not remove all axonal release, striatal microdialysis detects activity-independent extracellular dopamine, and nanofilm sensors also detect activity-independent axonal release in cultured dopamine neurons ^17,22,23^. This suggests that spontaneous dopamine release occurs in axons. Altogether, evoked and spontaneous dopamine release from axons, somata, and dendrites may all contribute to dopamine function and to its dysregulation in dopamine-associated disorders ^18,19,21,24–27^.

Here, we assessed roles of Munc13 in somatodendritic dopamine release. Munc13 is important for synaptic vesicle fusion in axons across species and cell types and acts as a vesicle priming protein ^28–36^. In mice with Munc13 ablation in dopamine neurons, we found a strong reduction in evoked somatodendritic dopamine release measured as D2-IPSCs. Dopamine release examined with a fluorescent sensor revealed a similar disruption when comparing somatodendritic release in the midbrain with axonal release in the striatum. Spontaneous dopamine transmission in the midbrain was unaffected by Munc13 ablation, bolstering the model of a separate secretory pathway. Finally, a small component of the midbrain D2-IPSC was due to norepinephrine innervation, reminiscent of the observation that D2 receptor-based dopamine sensors can report norepinephrine ^37^. The relative norepinephrine contribution to D2-IPSCs was increased after Munc13 ablation from dopamine neurons. Overall, these findings establish that Munc13 mediates evoked somatodendritic dopamine release. Our work supports that mechanisms for evoked somatodendritic release resemble those for axonal transmitter secretion and suggest the existence of pre-assembled release sites for efficient and temporally precise dopamine signaling.

## Results

### Munc13 ablation disrupts evoked somatodendritic dopamine release

Roles of Munc13 in somatodendritic dopamine release were first examined using whole-cell recordings in horizontal slices generated from mice lacking Munc13 proteins. We pursued a genetic strategy previously used to study axonal dopamine release ^35^ with dopamine neuron-specific ablation of Munc13-1 and constitutive knockout of Munc13-2 and Munc13-3 (Munc13 cKO^DA^, Fig. 1A). In whole-cell recordings from dopamine neurons (Fig. 1B) of Munc13 cKO^DA^ and Munc13 control mice, the spontaneous firing rates and inward currents induced by a hyperpolarizing step (I_H_) were similar (Fig. 1C-F). The input resistance (R_M_) and cell capacitance of these neurons were similar as well, but small magnitude decreases in R_M_ and increases in capacitance were detected (Fig. 1G+H). This is consistent with the modestly altered axon structure in the striatum of Munc13 cKO^DA^ mice ^35^. Midbrain dopamine transmission was measured via D2-IPSCs recorded in the SNc evoked by extracellular electrical stimulation (5 stimuli, 40 Hz) at varying intensities (Fig. 2A-C). At the lowest intensity (50 µA), neither Munc13 cKO^DA^ nor Munc13 control cells exhibited an appreciable D2-IPSC. As stimulus intensity increased, D2-IPSCs were induced in dopamine neurons of Munc13 control and Munc13 cKO^DA^ mice. D2-IPSC amplitudes in Munc13 cKO^DA^ slices were reduced by 73% at 150 µA stimulation intensity, 74% at 250 µA, and 67% at 350 µA. In both genotypes, D2-IPSCs were mediated by action potentials as they were eliminated by acute bath application of the Na^+^ channel blocker tetrodotoxin (TTX; Fig. 2D+E).

**Figure 1.**
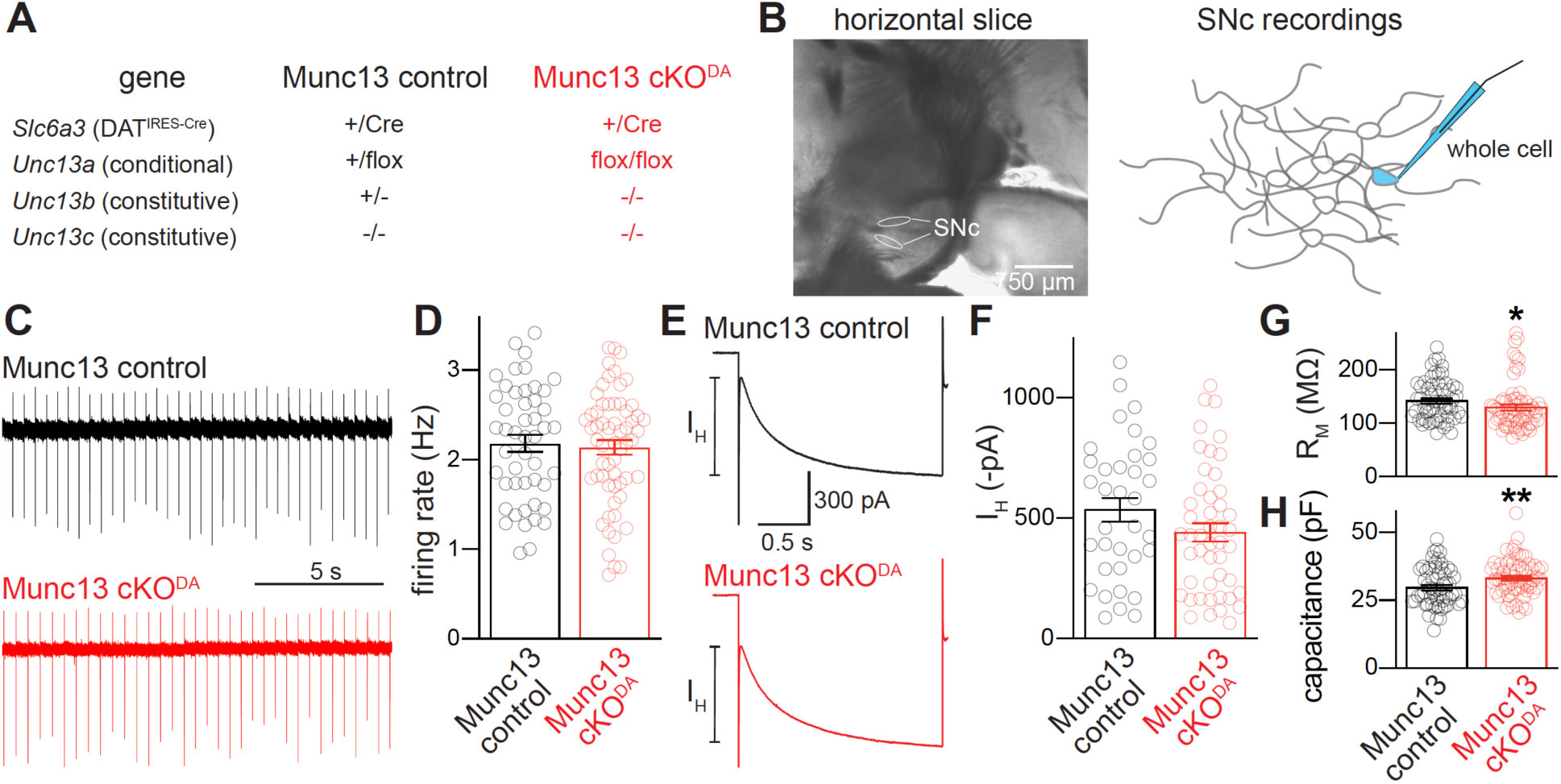
Electrical properties of dopamine neurons after Munc13 ablation. (A) The genetic strategy for Munc13 ablation as established before ^35^; Munc13-1 ablation is dopamine neuron-specific, Munc13-2 and Munc13-3 ablations are constitutive. (B) Brightfield image of a horizontal brain slice showing the substantia nigra pars compacta (SNc, left) and a schematic of the whole-cell recordings (right). (C, D) Example traces (C) and quantification of rates (D) of spontaneous firing of dopamine neurons, measured cell-attached over two to four minutes prior to break-in; Munc13 control 48 cells from 34 slices of 18 mice, Munc13 cKO^DA^ 60/40/21. (E, F) Example traces (E) and quantification of the amplitude (F) of HCN-mediated inward currents (I_H_) in response to a −50 mV, 2-s hyperpolarizing step; Munc13 control 35/25/15, Munc13 cKO^DA^ 47/32/15. (G, H) Quantification of the input resistance (G) and cell capacitance (H) measured after break-in; Munc13 control 57/33/16, Munc13 cKO^DA^ 64/45/15. Data are shown as mean ± SEM, * p < 0.05, ** p < 0.01 as assessed by unpaired t-tests (D, H), or Mann-Whitney rank sum tests (F, G).

**Figure 2.**
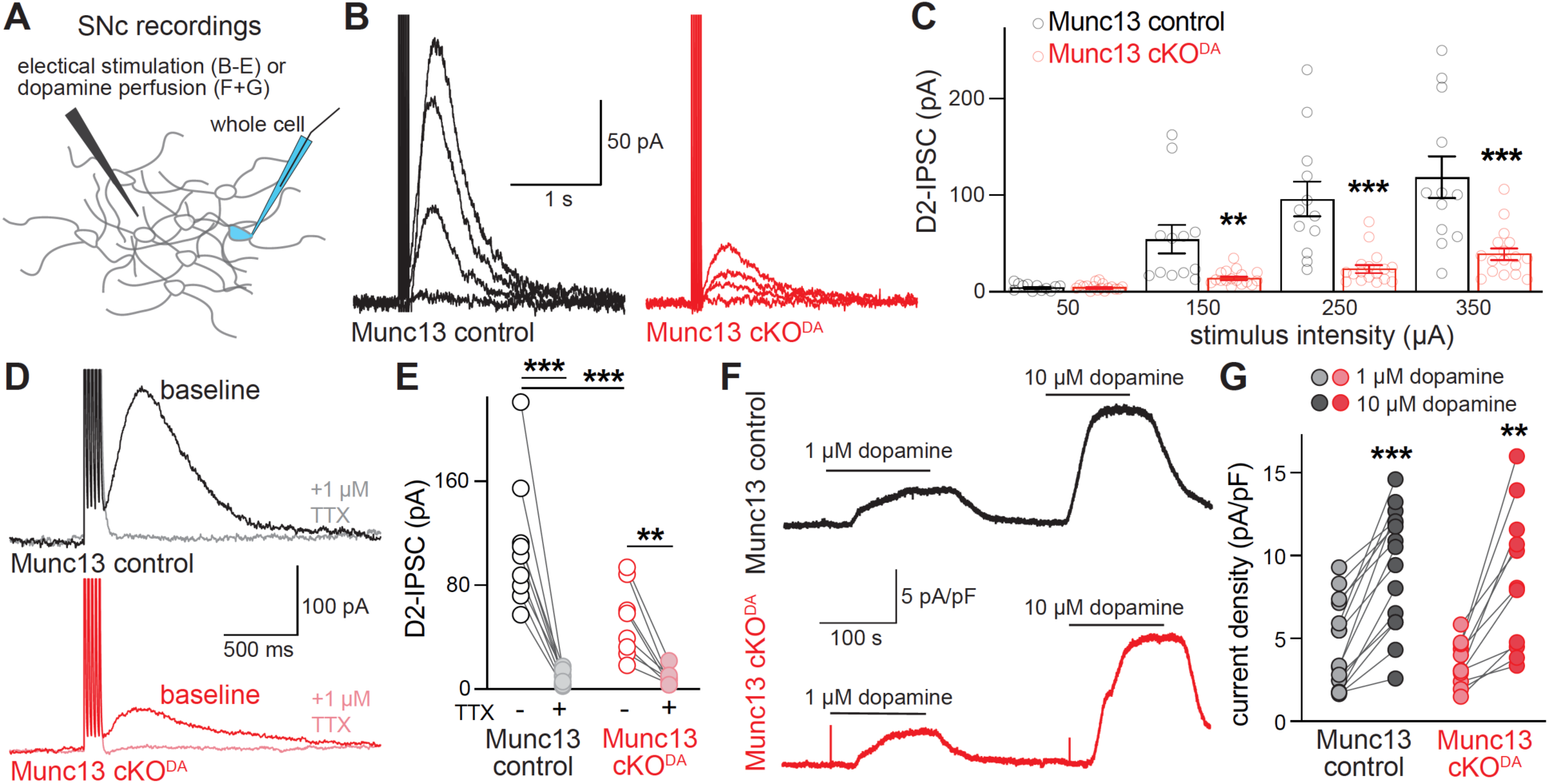
Munc13 ablation disrupts dopamine release in the SNc. (A) Schematic of D2-IPSC recordings in the SNc. (B, C) Example traces (B) and quantification of peak amplitudes (C) of D2-IPSCs induced by five stimuli at 40 Hz at 50, 150, 250, or 350 µA stimulation intensity; Munc13 control 12 cells from 10 slices of 6 mice, Munc13 cKO^DA^ 17/15/7. (D, E) Example traces (D) and quantification of peak amplitudes (E) of D2-IPSCs induced by five stimuli at 40 Hz at 350 µA stimulation intensity before or after perfusion of the voltage-gated Na^+^ channel blocker tetrodotoxin (TTX, 1 µm); Munc13 control 9/9/6, Munc13 cKO^DA^ 8/8/7. (F, G) Example traces (F) and quantification (G) of current density induced by 1 µM or 10 µM dopamine perfusion for ∼2 min until a peak current was achieved (calculated as the change in holding current divided by cell capacitance); Munc13 control 15/15/7, Munc13 cKO^DA^ 11/11/9. Data are shown as mean ± SEM, ** p < 0.01, *** p < 0.001 as assessed by two-way ANOVA in C (genotype: ***, intensity: ***, interaction: ***), E (genotype: *, TTX: ***, interaction: **) and G (genotype: ns, concentration: ***, interaction: ns) followed by Sidak’s multiple comparison tests (C, E, G, significance indicated in panels).

We next tested whether the genetic manipulations might alter D2 receptor activation. We measured D2 receptor-mediated currents induced in response to perfusion of 1 μM or 10 μM exogenous dopamine (Fig. 2F+G). Quantification of the current density did not reveal a difference between Munc13 cKO^DA^ and control neurons. This establishes that somatodendritic D2 receptors are present in Munc13 cKO^DA^ neurons and that these receptors detect exogenous dopamine similar to the control condition. These observations support that the reduced D2-IPSC amplitudes are a result of decreased dopamine release from the somatodendritic compartment following ablation of Munc13.

### Munc13 ablation similarly disrupts axonal and somatodendritic release

A more direct measure of dopamine release was next obtained by imaging fluorescence changes of GPCR-activation based dopamine (GRAB_DA_) sensors. In a first set of imaging experiments, we used widefield microscopy as established before to assess axonal dopamine release in the striatum with the D2 receptor-based dopamine sensor GRAB_DA2m_ ^24,38^ (Fig. 3A). GRAB_DA_ was expressed with AAVs injected into the striatum. Acute brain slices were prepared several weeks after injection and GRAB_DA_ fluorescence changes were measured and quantified as ΔF/F_0_ in a large field of view. GRAB_DA_ fluorescence transients in response to either single stimuli or trains of 10 stimuli at 10 Hz (Fig. 3B-G) were strongly reduced in Munc13 cKO^DA^ slices. This confirms previous work that defined Munc13 proteins as important mediators of evoked axonal dopamine release with carbon fiber amperometry in acute striatal brain slices ^35^.

**Figure 3.**
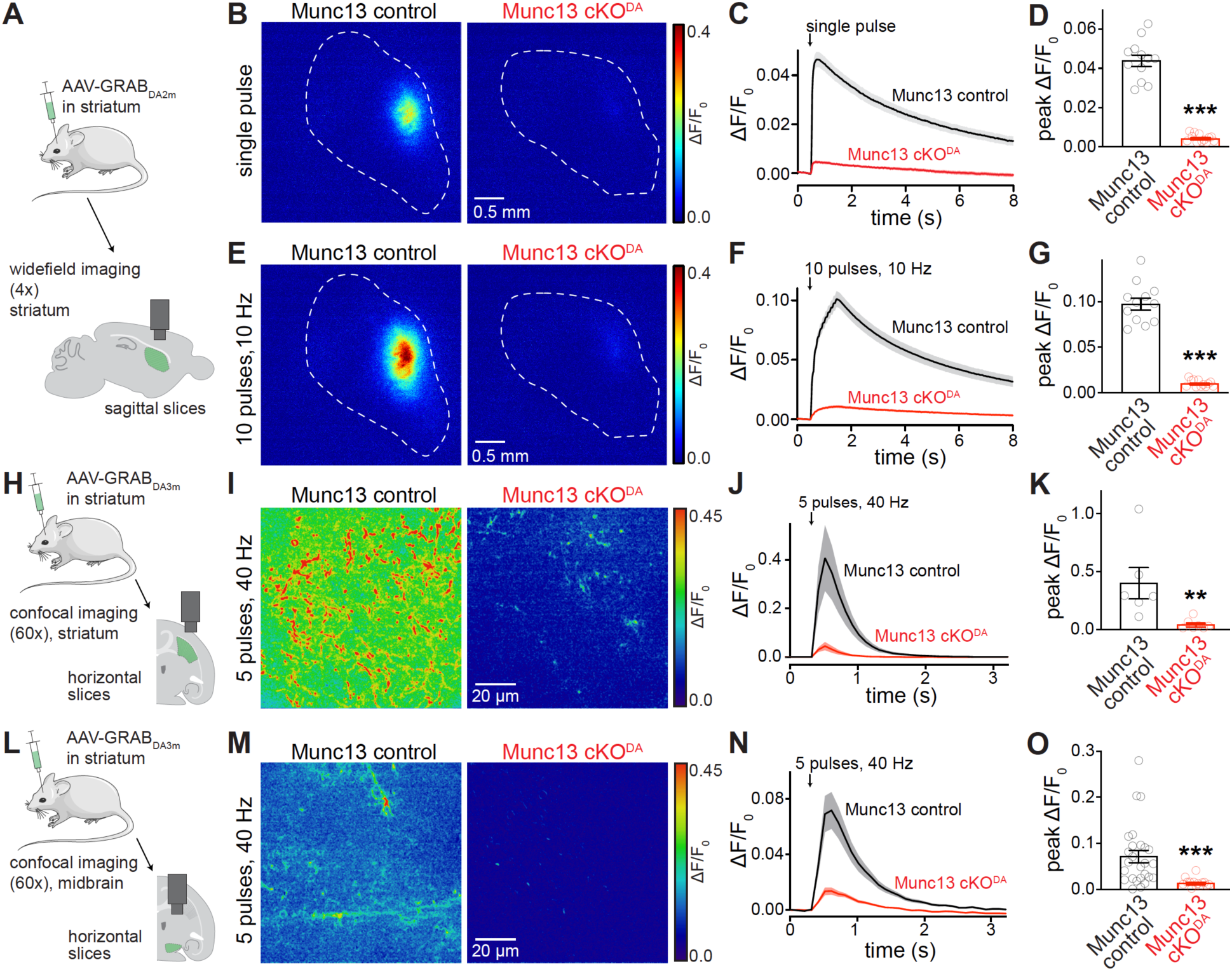
Munc13 ablation disrupts striatal and midbrain dopamine release monitored with GRAB_DA_. (A) Schematic of striatal unilateral AAV injections followed by widefield imaging in parasagittal striatal slices. (B-D) Example ΔF/F_0_ images (B) and quantification of the of ΔF/F_0_ time course (C) and peak amplitude (D) during single extracellular electrical stimuli; Munc13 control 12 images of 12 slices from 3 mice, Munc13 cKO^DA^ 12/12/3. (E-G) As in B-D, but for 10 stimuli at 10 Hz; Munc13 control 12/12/3, Munc13 cKO^DA^ 12/12/3. (H) Schematic of bilateral striatal AAV injections followed by confocal imaging in hemisected horizontal striatal slices. (I-K) Example ΔF/F_0_ images (I) and quantification of the of ΔF/F_0_ time course (J) and peak amplitude (K) during five electrical stimuli at 40 Hz at 250 μA stimulation intensity; Munc13 control 6/4/4, Munc13 cKO^DA^ 7/3/3. (L) Schematic of bilateral striatal AAV injections followed by confocal imaging in hemisected horizontal midbrain slices. (M-O) As in I-K, but for horizontal midbrain slices; Munc13 control 26/6/3, Munc13 cKO^DA^ 14/6/4. Data are shown as mean ± SEM, ** p< 0.01, *** p < 0.001 as assessed by unpaired t-tests (D, G) or Mann-Whitney rank sum test (K, O).

In a next set of experiments, we used a retrograde AAV strategy expressing GRAB_DA3m_, a D1-receptor based dopamine sensor with an increased signal-to-noise ratio compared to GRAB_DA2m_ ^39^. We injected AAV-GRAB_DA_ into the striatum to transduce axonal collaterals of striatal neurons and neurons with afferent projections into the striatum. This strategy transduces dopamine neurons via their axons without disrupting their somatodendritic compartment in the midbrain. GRAB_DA_ fluorescence was then imaged using spinning-disc confocal microscopy in smaller fields of view compared to the wide-field imaging experiments described above and with a stimulus protocol (5 stimuli at 40 Hz) well-suited for midbrain measurements. We used this approach in parallel in the striatum and the midbrain to assess dopamine release in both areas (Fig. 3H+L). Similar to the wide-field measurements (Fig. 3A-D), there was a robust impairment of dopamine release in Munc13 cKO^DA^ slices in the striatum (Fig. 3H-K). The assessment of stimulus-induced GRAB_DA_ transients in the ventral midbrain of Munc13 cKO^DA^ and Munc13 control slices revealed a strong reduction in somatodendritic dopamine release (Fig. 3M-O). Overall, the reduction in somatodendritic dopamine release monitored with GRAB_DA_ was similar in magnitude to effects observed in striatal GRAB_DA_ experiments (Fig. 3A-H) and in midbrain D2-IPSC recordings (Fig. 2A-C). Hence, axonal and somatodendritic dopamine release have a comparable dependency on Munc13. The findings are reminiscent of previous work that established shared dependency of axonal and somatodendritic dopamine release on RIM active zone scaffolds and on the Ca^2+^ sensor Synaptotagmin-1 ^12,17,18,22,35,40^.

### Spontaneous somatodendritic dopamine transmission persists after Munc13 cKO^DA^

Unlike evoked release, spontaneous somatodendritic dopamine release is insensitive to blockade of voltage-gated Na^+^ channels, voltage-gated Ca^2+^ channels, lowering of extracellular Ca^2+^, or genetic ablation of RIM or Synaptotagmin-1 from dopamine neurons ^10,12,18,21^. Spontaneous glutamate or GABA release at fast synapses is greatly reduced when Munc13 is ablated ^31,41^. We examined whether spontaneous dopamine release shows a similar dependency on Munc13, assessing slices of Munc13 cKO^DA^ and Munc13 control mice. We used a previously established strategy to facilitate the detection of spontaneous D2-IPSCs by exogenous expression of D2 receptors via AAVs injected into the midbrain (Fig. 4A) ^12,21^. In whole cell recordings in acute brain slices approximately two weeks after AAV transduction, neither the amplitude nor the frequency of spontaneous D2-IPSCs were changed in Munc13 cKO^DA^ slices (Fig. 4B-D). To assess whether the D2 receptor overexpression strategy altered the phenotypes observed in response to electrical stimulation, evoked D2-IPSCs were measured. Similar to the D2-IPSCs mediated by endogenous receptors (Fig. 2), the current amplitudes were severely reduced after ablation of Munc13 in slices expressing exogenous D2 receptors (Fig. 4E+F). Hence, expression of exogenous D2 receptors did not alter the impairment in stimulated somatodendritic dopamine release. Previous work describing the independence of spontaneous release on RIM or Synaptotagmin-1, or on Ca^2+^ entry, led to the model that spontaneous release in the midbrain occurs through a secretory pathway that is distinct from that for evoked release ^12,18,21^. The lack of reliance on Munc13 strengthens the model of a separate release pathway for spontaneous release in the midbrain.

**Figure 4.**
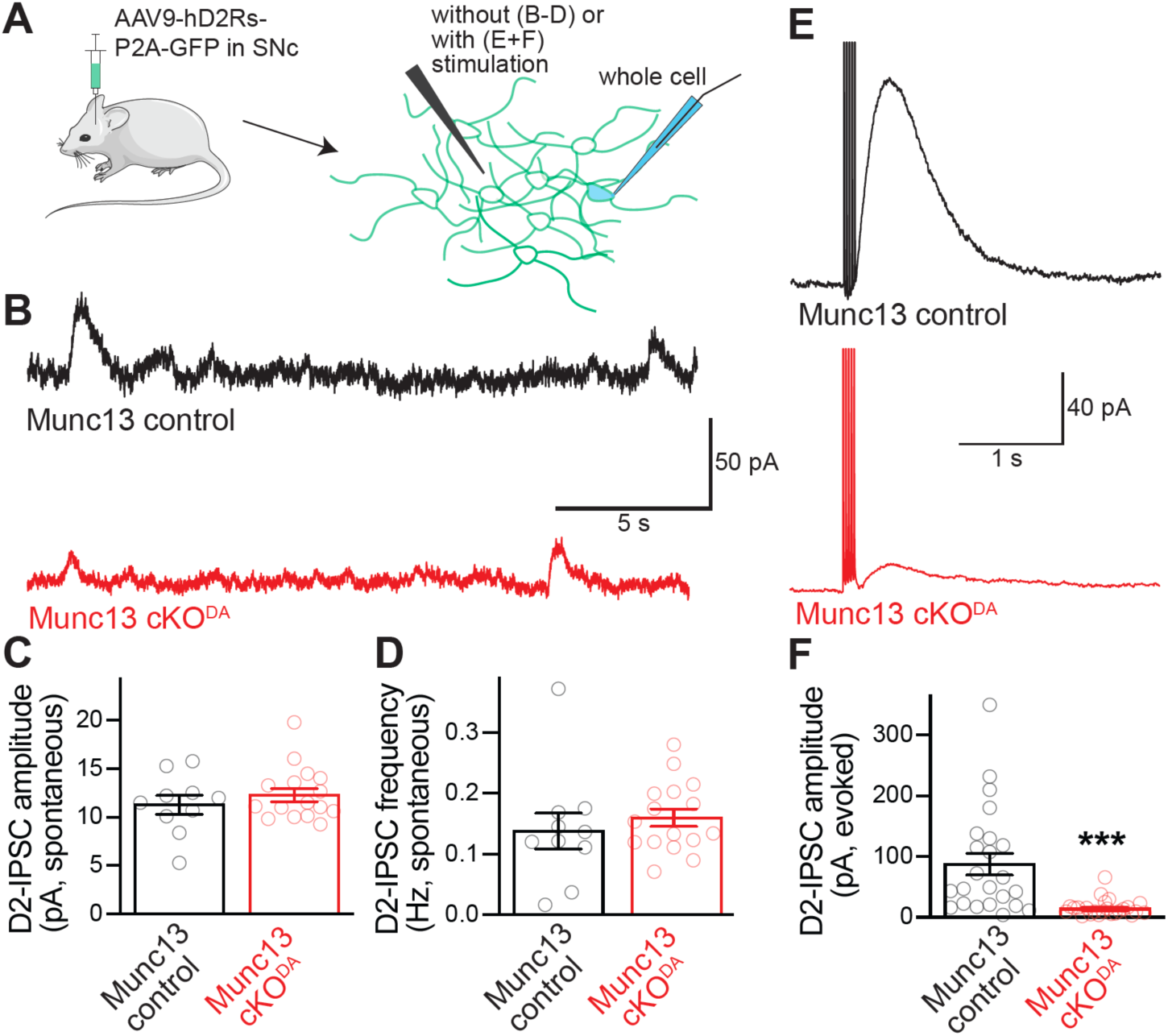
Spontaneous dopamine release in the SNc persists after Munc13 ablation. (A) Schematic of AAV-mediated D2R expression followed by whole-cell recordings of SNc dopamine neurons. (B-D) Example traces (B) and quantification of amplitude (C) and frequency (D) of spontaneous D2-IPSCs; Munc13 control 10 cells from 4 slices of 4 mice, Munc13 cKO^DA^ 16/8/4. (E, F) Example traces (D) and quantification of peak amplitudes (E) of D2-IPSCs induced by five stimuli at 40 Hz and 150 µA stimulation intensity; Munc13 control 23/17/4, Munc13 cKO^DA^ 24/13/4. Data are shown as mean ± SEM, *** p < 0.001, as assessed by Mann-Whitney rank sum tests (C, F) or an unpaired t-test (D).

### Norepinephrine innervation modestly contributes to D2-IPSCs measured in the midbrain

The residual component of dopamine release in Munc13 cKO^DA^ mice could be explained by Munc13 independence, by the presence of a small amount of a left-over Munc13 protein, or by neurotransmitters other than dopamine that might activate D2 receptors or dopamine sensors ^35,37,41^. The substantia nigra receives norepinephrine innervation, and neurotransmitter released from these axons might act on D2 receptors ^37,42–44^. To examine this possibility, D2-IPSCs were measured before and after inhibition of release from norepinephrine terminals by the selective α2-adrenergic receptor agonist UK14,304 (Fig. 5A). Activation of this G_i_-coupled autoreceptor is known to inhibit stimulation-evoked norepinephrine release ^45–47^. D2-IPSCs recorded in control slices showed a modest (17%) but statistically significant decrease in amplitude after bath addition of UK14,304, which was reversed by the α2 adrenergic receptor antagonist idazoxan (Fig. 5B-E). In neurons recorded in Munc13 cKO^DA^ slices, the remaining D2-IPSC was reduced by a similar absolute magnitude in response to bath application of UK14,304, though the smaller initial D2-IPSC amplitude before application of UK14,304 resulted in a larger relative inhibition (47%), which was also reversed by idazoxan (Fig. 5B-E). We conclude that only a small component of the D2-IPSC is mediated by norepinephrine axons. While this mechanism contributes significantly to the measured D2-IPSC that remains after Munc13 ablation, it does not fully account for it, and a small amount of somatodendritic dopamine release persists after the genetic ablation of Munc13 tested here.

**Figure 5.**
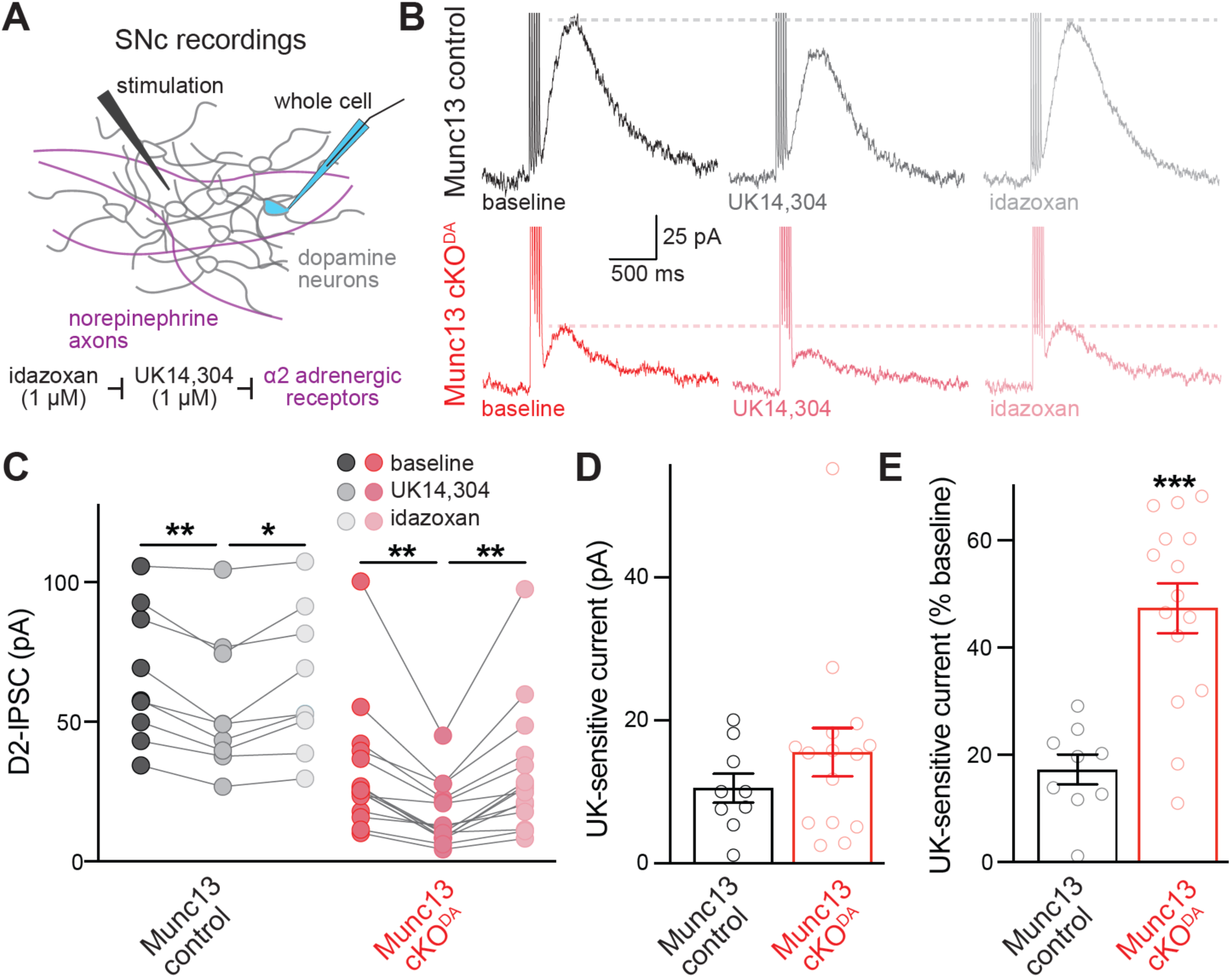
Contribution of release from norepinephrine axons to D2-IPSCs in the SNc. (A) Schematic and pharmacological strategy of D2-IPSC recordings in the SNc. (B-E) Example traces (B) and quantification of peak amplitudes (C) of D2-IPSCs induced by five stimuli at 40 Hz and 250 μA stimulation intensity, and of absolute (D) and normalized (E) UK14,304 sensitivity in an experiment with sequential addition of UK14,304 and idazoxan; Munc13 control 9 cells from 9 slices of 4 mice, Munc13 cKO^DA^ 15/15/6. Data are shown as mean ± SEM, * p < 0.05, ** p < 0.01, *** p < 0.001 as assessed by a two-way ANOVA (C; genotype: ***, drug: ***, interaction: ns) followed by Tukey’s post tests (indicated in figure), a Mann-Whitney rank sum test (D), or an unpaired t-test (E).

## Discussion

The presented D2-IPSC recordings and GPCR-based sensor measurements establish that the vesicle priming protein Munc13 is important for evoked somatodendritic dopamine release. Effect magnitudes of Munc13 ablation are similar for axonal and somatodendritic dopamine release. There is a small midbrain D2-IPSC remaining after Munc13 ablation, and about half of it is accounted for by release from norepinephrine axons. Contrasting evoked somatodendritic release, spontaneous D2-IPSCs are independent of Munc13 in dopamine neurons. Overall, our work strengthens the model of pre-organized release sites for rapid and precise somatodendritic dopamine secretion.

### Are there preorganized sites to prime vesicles for somatodendritic dopamine release?

For spatiotemporally precise secretion in response to action potentials, fusion-ready vesicles must be tethered close to their future sites of release ^48,49^. In addition, voltage-gated Ca^2+^ channels must be concentrated near the fast vesicular Ca^2+^ sensors ^2,50,51^. At conventional synapses and in dopamine axons, active zone proteins orchestrate these processes for release to be initiated within a millisecond of an arriving action potential ^52,53^.

Whether similar protein assemblies mediate somatodendritic neurotransmitter secretion has remained an open question. Munc13’s importance provides molecular evidence for vesicle priming in somatodendritic dopamine secretion. Munc13 is the principal priming protein at synapses, where it catalyzes the assembly of SNARE complexes ^30,31,54–56^. Munc13 also supports the tight docking of synaptic vesicles ^32,33,57^. Central to the roles of Munc13 in these processes is its recruitment and monomerization by RIM, active zone scaffolds that activate Munc13 ^33,54,58–60^.

The findings that Munc13 (Figs. 2+3) and RIM ^18^ are important for stimulus-induced somatodendritic dopamine release support that the synaptic vesicle priming mechanism operates in somatodendritic dopamine secretion. More broadly, these molecular requirements indicate that somatodendritic secretion occurs at pre-assembled exocytic sites at the plasma membrane of somata and/or dendrites and imply that the vesicles are docked before fusion. While it has remained difficult to morphologically define somatodendritic release sites, the data presented here and in previous studies ^10,12,18,61^ provide a foundation to identify them in the future. They predict the clustered occurrence of RIM and Munc13 at the somatodendritic plasma membrane, and the presence of a vesicular compartment containing the Synaptotagmin-1. Because these proteins typically mediate the fusion of synaptic vesicles ^2,52,62^, these findings also suggest that somatodendritic release is mediated by small, clear vesicles. This model differs from early studies that suggested fusion of tubulovesicular structures for somatodendritic release and is more consistent with reports on the presence of small, pleiomorphic vesicles ^63–66^. Overall, this and previous work identifies molecules important for somatodendritic dopamine release.

### Organization of somatodendritic transmission beyond vesicle priming

Many features of somatodendritic transmission remain understudied. While previous work indicates tight coupling of Ca^2+^ entry and secretion with a high initial release probability, the Ca^2+^ sources and their targeting to release sites remain uncertain ^19,67,68^. Deletion of the fast Ca^2+^ sensor Synaptotagmin-1 strongly impairs single pulse D2-IPSCs, but stimulus trains evoke significant somatodendritic release in Synaptotagmin-1 knockout dopamine neurons ^12^. Knockdown, knockout, and antibody application experiments have also implicated Synaptotagmin-4 and Synaptotagmin-7 ^13,14,16,69^. While vertebrate Synaptotagmin-4 is unlikely to mediate Ca^2+^ sensing ^70^, effects of Synaptotagmin-7 ablation or inhibition are mild. Overall, the full complement of Ca^2+^ sensors for somatodendritic dopamine release remains unknown. A determinant for efficacy of signaling is the positioning of receptors relative to release sites. At synapses, tight coupling at tens of nanometers controls receptor activation ^62^. For somatodendritic dopamine transmission, release-receptor organization remains unclear. Morphological studies have identified D2 receptor clusters ^71–73^, but it has not been possible to assess their proximity to release sites. Functional imaging has established that somatodendritic dopamine release generates hotspots, and that uncaging a D2 receptor antagonist inhibits receptor activation within the first ∼100 ms of stimulation, but not at later time points ^74^. In addition, D2-IPSC induction relies on a high dopamine concentration, 30 – 100 μM ^61^. These studies suggest that dopamine acts rapidly and in a contained space. It is noteworthy, though, that dopamine diffusion is fast and receptors several μm away from release sites might be reached within 100 ms ^75,76^. Future studies should assess release-receptor organization in the midbrain dopamine system to define dopamine’s signaling mechanisms.

### Norepinephrine contribution to somatodendritic dopamine signaling

Norepinephrine neurons innervate the midbrain ^44,77,78^, and α1 adrenoreceptors inhibit dopamine neuron activity ^79^. Norepinephrine may also act as an agonist of D2 and D2-like receptors ^42,80^, and it activates D2-receptor based dopamine sensors ^37^. Here, we find that norepinephrine innervation contributes to D2-IPSCs, boosting this inhibition. In control mice, the contribution is small (Fig. 5). In Munc13 cKO^DA^ mice, norepinephrine innervation accounts for half of the remaining D2-IPSC. Whether released norepinephrine activates D2 receptors, or whether norepinephrine neurons might co-release dopamine before its conversion to norepinephrine ^20,81,82^, remains unclear.

### Spontaneous somatodendritic dopamine release

Dopamine is associated with movement initiation and vigor, but how dopamine controls movement remains debated ^24,83–88^. Recent studies suggest that spontaneous release suffices to mediate movement. Mice that lack RIM in dopamine neurons have disrupted in vivo dopamine dynamics. In these mice, movement initiation is unaffected, but dopamine depletion or receptor blockade disrupts their movement ^24^. Spontaneous somatodendritic dopamine release is unaffected by RIM ablation ^18^. The work presented here bolsters the conclusion that evoked dopamine release is dispensable for movement. Munc13 cKO^DA^ mice have a large reduction in evoked dopamine release (Figs. 2+3). While their body weight at weaning is reduced, they survive, move, and eat ^35^.

The cellular pathway for spontaneous somatodendritic dopamine release remains to be defined. Previous work established that it is vesicular, but independent of voltage-gated Na^+^ channels, voltage-gated Ca^2+^ channels, extracellular Ca^2+^, intracellular Ca^2+^ stores, and Synaptotagmin-1 expression in dopamine neurons ^12,21^. Here, we show that spontaneous somatodendritic transmission persist after Munc13 ablation from dopamine neurons (Fig. 4), in addition to its independence of RIM ^18^. The spontaneous dopamine release pathway may not only be different from that of evoked release, but it might also differ from that of spontaneous release at synapses, where RIM, Munc13, and Synaptotagmin-1 contribute across synapses, cell types and species ^31,41,59,89–91^. What could those pathways be? Distinct SNARE proteins or their Ca^2+^-dependent regulators may control spontaneous release, and some proteins might specifically control spontaneous fusion, for example Vti1a, Vamp7, or Doc2 ^92–94^. Conversely, ablation of the canonical SNAREs synaptobrevin/VAMP-2, Syntaxin-1 and SNAP25 strongly decreases spontaneous release at synapses, indicating that most spontaneous release relies on them and shares vesicles with evoked release ^2^. It is possible that the feature of distinct vesicle pools or SNARE proteins is prominent in dopamine neurons. Recent work has identified specific adaptor complexes for vesicles released during high-frequency firing of dopamine neurons ^95^. Similar mechanisms could account for a pool of spontaneous vesicles. Alternatively, spontaneous dopamine release may rely on a different pathway or be mediated by different cells. For the former, the exocyst is a candidate pathway that consists of conserved secretory machinery, and it typically participates in neuronal development but not transmitter exocytosis ^96,97^.

Overall, our studies provide a molecular handle on the identity of the sites that underlie evoked release in the midbrain, as they indicate that Munc13 might be marker for these sites ^98,99^. The alternate pathways described above should be systematically investigated for a role in spontaneous somatodendritic dopamine release.

## Acknowledgements

We thank members of the Williams and Kaeser laboratories for insightful discussions and feedback, and R. López and K. Nozawa for comments on the manuscript. This work was supported in part by the NIH (R01DA056109, JTW and PSK; R01NS103484, R01NS083898, PSK; R01DA004523, T32DA007262, JTW) and Harvard Medical School (PSK). We acknowledge N. Brose and C. Imig for sharing Munc13-1, -2 and -3 mutant alleles, and Y. Li for providing access to GRAB_DA_ AAVs. AB is a recipient of a Brain and Behavior Research Foundation Young Investigator Grant (#31271). The mouse images in Figs. 3 and 4 were provided by Servier Medical Art (https://smart.servier.com) licensed under CC BY 4.0 (https://creativecommons.org/licenses/by/4.0/).

## Author contributions

Conceptualization, JJL, AB, JTW and PSK; Methodology, JJL, AB, GH and JTW; Formal Analysis, JJL, AB, JTW and PSK; Investigation, JJL, AB and JTW; Resources, AB and GH; Writing-Original Draft, JJL and PSK; Writing-Review & Editing, JJL, AB, GH, JTW and PSK; Supervision, JTW and PSK; Funding Acquisition, JTW and PSK.

## Declaration of interests

The authors declare no competing interests.

## Materials and Methods

### Mice

Munc13 cKO^DA^ mice were crossed as described before ^35^. Conditional Munc13-1 floxed mice (MGI:7276178) ^35^ were crossed to constitutive Munc13-2 knockout mice (MGI:2449706) ^31^, constitutive Munc13-3 knockout mice (MGI:2449467) ^100^, and DAT^IRES-cre^ mice (JAX: 006660) ^101^. Munc13 cKO^DA^ mice were homozygote for Munc13-1 floxed, Munc13-2 null, and Munc13-3 null, and heterozygote for DAT^IRES-cre^. Munc13 control mice were heterozygote for Munc13-1 floxed and Munc13-2 null, homozygote for Munc13-3 null, and heterozygote for DAT^IRES-Cre^. We used littermate mice or age-matched mice from the same breeding colony for experiments. Mice were genotyped in the lab or by Transnetyx. Experiments were performed in accordance with protocols approved by Animal Care and Use committees at Harvard Medical School and Oregon Health and Science University.

### Brain slice preparation

For D2-IPSC recordings and GRAB_DA3m_ imaging, mice (110 to 209 days old) were deeply anesthetized with isoflurane and decapitated. The brain was dissected into warm (32 to 35° C) Krebs buffer containing (in mM): 126 NaCl, 2.5 KCl, 1.2 MgCl_2_, 2.4 CaCl_2_, 1.4 NaH_2_PO_4_, 25 NaHCO_3_, and 11 dextrose. Krebs buffer was bubbled with 95% O_2_/5% CO_2_, pH 7.4, prior to decapitation and continuously throughout cutting and recovery and contained 10 µM MK-801. The brain was then sliced on a vibrating microtome. Horizontal brain slices containing the ventral midbrain were cut at a thickness of 222 µm and allowed to recover for at least 30 min at 30 to 32° C prior to recording or imaging. For GRAB_DA2m_ imaging, mice (84 to 127 days old) were deeply anesthetized with isoflurane and decapitated. Brains were dissected out and 250 μm thick parasagittal sections of the striatum were cut on a vibrating microtome in ice cold cutting solution containing (in mM): 7.5 MgSO_4_, 75 sucrose, 75 NaCl, 1 NaH_2_PO_4_, 12 glucose, 26.2 NaHCO_3_, 2.5 KCl, 1 sodium ascorbate, 1 myo-inositol, and 3 sodium pyruvate bubbled with 95% O_2_/5% CO_2_, pH 7.4, 300 to 305 mOsm. Slices were incubated in recovery solution for at least 1 hour at room temperature (21 to 25° C) containing (in mM): 126 NaCl, 1 NaH_2_PO_4_, 2.5 KCl, 2 CaCl_2_, 12 glucose, 1.3 MgSO4, 12 glucose, 26.2 NaHCO3, 1 sodium ascorbate, 1 myo-inositol, and 3 sodium pyruvate bubbled with 95% O_2_/5% CO_2_, pH 7.4, 300 to 305 mOsm.

### Electrophysiology

Experiments were performed using previously established methodology ^10,12,18,21^. Slices were bisected along the midline and experiments were carried out in hemi-sections placed in a recording chamber under continuous perfusion with Krebs buffer (2 to 3 ml/min) at 34° C bubbled with 95% O_2_/5% CO_2_, pH 7.4. Krebs buffer for recording also contained NBQX (600 nM), CGP55845 (300 nM), and picrotoxin (100 µM). In hemi-sected, horizontal midbrain slices, SNc dopamine neurons were identified by size, morphology, and position lateral to the medial terminal nucleus of the accessory optic system. Recordings were made using glass pipettes (initial resistance: 1.2 to 1.6 MΩ) filled with an internal solution containing (in mM): 100 K-methanesulfonate, 20 NaCl, 1.5 MgCl_2_, 10 HEPES (K), 2 ATP, 0.3 GTP, 10 phosphocreatine, and10 BAPTA (4K), pH 7.4, 270 to 290 mOsm. Firing rates were recorded for ≥ 1 min in cell attached mode prior to break-in and action potentials were identified using an amplitude threshold. After break-in, the holding potential was set to -55 mV and cell capacitance, membrane resistance, and series resistance were measured with a 5 mV test pulse. The uncompensated series resistance was monitored and cells with an initial series resistance ≥ 10 MΩ were excluded from experiments. The I_h_ current was measured as the average amplitude over the last 25 ms of a 2 s, -50 mV hyperpolarizing voltage step. D2-IPSCs were recorded no sooner than 5 min after break-in. A monopolar electrode was positioned approximately 75 to 100 µm caudal to the recorded cell and D2-IPSCs were induced with a train of five stimuli at 40 Hz using 50 µA, 150 µA, 250 µA, and 350 µA stimulus intensities. The order of stimulation (ascending or descending) was alternated between cells. Each intensity was tested with three to five consecutive sweeps with an inter-sweep interval of one minute and the average amplitude of those sweeps was quantified at each stimulus intensity. Series resistance was monitored between different stimulus intensities and cells with a series resistance that exceeded 10 MΩ were discarded. The sensitivity to voltage-gated Na^+^ channel blockade was examined by comparing the D2-IPSC amplitude evoked with a 350 µA stimulus before and after the addition of 1 µM TTX. Spontaneous D2-IPSCs were measured over a five-minute period starting no sooner than 10 min after break-in and detected automatically using a sliding template in Axograph. In a second analyses, manual identification of spontaneous D2-IPSCs by an experimenter blind to genotype yielded results that were similar to those measured with the template function and shown in Fig. 4C+D. D2 receptor activation by exogenous dopamine was measured by superfusing dopamine (1 µM or 10 µM) and is reported as the maximum change in current density. First, 1 µM dopamine was superfused until a peak current was attained (for ∼2 min) and then removed. After the current returned to baseline, 10 µM dopamine was superfused until a new peak was attained (for ∼2 min). Current density was calculated by dividing the amplitude of the change in holding current by the recorded cell’s capacitance. Norepinephrine axon contribution was estimated by sequential perfusion of UK14,304 (1 µM) followed by idazoxan (1 µM). Effects were calculated using the final three sweeps (at 3 to 5 min) during an 8 min perfusion for each condition. Representative traces of evoked D2-IPSCs are the average of three consecutive sweeps. Representative traces of pacemaker firing, dopamine superfusion, and spontaneous D2-IPSCs are from single sweeps.

### Stereotaxic surgeries

GRAB_DA_ and D2R were delivered via stereotaxic injections of AAVs into the mouse brain. AAVs for D2R expression were custom made as described ^12^, and AAVs for GRAB_DA_ expression were purchased (in some cases with permission from Y. Li). For AAV9-GRAB_DA2m_ ^38^ (also called AAV-hSyn-GRABDA2m, rAAV-hSyn-DA4.4; serotype: AAV2/9; Biohippo BHV12400441-9), 36 to 67 day-old mice were anesthetized with isoflurane and mounted on a stereotaxic frame. Anesthesia was maintained throughout surgery using isoflurane. After exposure of the skull, a burr hole was drilled for a unilateral AAV injection. Next, 1 μl of AAV (1 to 2 x 10^12^ viral genomic copies/ml) was injected into the dorsal striatum a rate of 0.1 µl/min using a microinjection syringe pump, coordinates were: 1.0 mm anterior to Bregma, 2.0 mm lateral, 2.5 mm below pia. Post-operative analgesia was provided following approved protocols. Experiments were conducted 48 to 60 days following injection. For AAV-GRAB-DA_3m_ ^39^ (also called rAAV-hSyn-DA3m-WPRE-PA; serotype: AAV2/RETRO, Biohippo BHV12400547-12), 70 to 150 day-old mice were anesthetized with isoflurane and mounted on a stereotaxic frame. Anesthesia was maintained throughout surgery using isoflurane. After exposure of the skull, two burr holes were drilled for bilateral AAV injections. On each side, 120 nl of AAV (titer: 5.0 x 10^12^ viral genomic copies/ml) were injected into the striatum at three locations along the dorsal/ventral axis. Virus was injected at a rate of 0.3 µl/min using a microinjection syringe pump, coordinates were: 1.25 mm anterior to Bregma, 1.50 mm lateral, and 3.40 mm, 3.30 mm, 3.20 mm below pia. Post-operative analgesia was provided following approved protocols. Experiments were conducted 20 to 45 days following injection. For AAV-hD2Rs-P2A-GFP ^12^ (custom made from pFB-AAV9-CAG-hD2Rshort-SNAP-p2A-GFP-WPRE-SV40pA, lab plasmid code p1038), 90 to 140 day-old mice were anesthetized with isoflurane and mounted on a stereotaxic frame. Anesthesia was maintained throughout surgery using isoflurane. After exposure of the skull, two burr holes were drilled for bilateral AAV injections. On each side,150 nl of AAV (titer: 2.15 x 10^13^ viral genomic copies/ml) were injected into the midbrain. Virus was injected at a rate of 0.3 µl/min using a microinjection syringe pump, coordinates were: 2.3 mm posterior to Bregma, 1.3 mm lateral, and 4.5 mm below pia. Analgesia was provided following approved protocols. Experiments were conducted 14 to 21 days after injection.

### Wide-field imaging in striatum

GRAB_DA2m_ imaging in parasagittal slices containing the dorsal striatum was performed following established methodology ^24,102^ with modifications. Images were acquired with a widefield fluorescence microscope (Olympus BX51) with a 470 nm excitation LED, a 4x objective and a scientific metal-oxide-semiconductor camera (sCMOS, Hamamatsu ORCA-Flash4.0) in artificial cerebrospinal fluid (ACSF) containing (in mM): 126 NaCl, 26.2 NaHCO_3_, 2.5 KCl, 1.3 MgSO_4_, 2 CaCl_2_, 1 NaH_2_PO_4_, and 12 glucose. The ACSF was heated to 34 to 36 °C and slices were perfused with a flow rate of at 2 to 3 ml/min, and 1 μM DhβE was included in the bath. Solutions were continuously bubbled with 95% O_2_/5% CO_2_, and recordings were completed within 5 h of slicing. Electrical stimulation was delivered with an intensity of 90 μA (biphasic wave, 0.25 ms in each phase) and with a linear stimulus isolator (A395, World Precision Instruments) through a unipolar glass pipette (tip diameter of 2 to 4 μm) filled with ACSF, using single pulses or 10 pulse stimulus trains at 10 Hz. Image acquisition, LED illumination and timed electrical stimulation was controlled with a digitizer (Molecular Devices, Digidata 1440A). LED power was kept constant at 10% across genotypes and repeats. Images were acquired at 512 × 512 pixels per frame, with a sampling frequency of 50 frames per second and an exposure time of 20 ms per frame. Fields of view were selected based on the presence of similar visible baseline GRAB_DA_ fluorescence prior to stimulation. Three sweeps were performed and averaged for each area for single stimuli and for 10 stimuli at 10 Hz. In this setup, a single pixel represents a striatal area of 6.4 × 6.4 μm^2^. For analyses, ImageJ/Fiji ^103^ was used and background fluorescence was estimated from regions of no sensor expression and subtracted from each image frame of the video. The pixels with intensity values in the top 50% of the intensity histogram of the image stack generated from background subtracted images was used for plotting the ΔF/F_0_ time course. F_0_ was calculated as the average GRAB_DA2m_ signal over 0.5 s (25 frames) immediately before stimulation (F_0_ in Munc13 control: 696.2 ± 92.7 arbitrary fluorescence units, 12 images/12 slices/3 mice, Munc13 cKO^DA^: 974.2 ± 131.7, 12/12/3, p = 0.09 as assessed by an unpaired t-test). ΔF/F_0_ was calculated for each pixel and quantified. For the peak plots, the maximum value of ΔF/F_0_ within 0.7 s or 1.1 s for single or train stimuli, respectively, was plotted. Sample heatmaps show the peak ΔF/F_0_ image frame after background subtraction and with brightness and contrast adjustments identical across genotypes; the display color range was set to illustrate the full range of the ΔF/F_0_ signal and the underlying raw ΔF/F_0_ was not saturated.

### Confocal imaging in striatum and midbrain

GRAB_DA3m_ imaging on hemisected horizontal slices was carried following methodology described before ^104^ but with an upright microscope (Olympus BX51 WI), a 60x objective (Olympus LUMPlanFL 60X/1.00 W) and a Yokogawa CSU10-B-F300-E-B488 spinning-disc confocal unit. Excitation was at 488 nm with a Cobolt 06-MLD laser controlled by a NEOS AOTF module. Fluorescence was detected and recorded at 10 Hz with a Photometrics Prime 95B camera. Hardware and acquisition settings were controlled by µManager2.0. Stimulation was carried out using Axograph and triggered by µManager2.0 at the start of each video to synchronize acquisition and stimulus timing. Stimulation was delivered with a monopolar electrode as described for evoked D2-IPSCs at 250 µA stimulation intensity, and 5 pulses at 40 Hz were applied. Fields of view were selected based on the presence of similar visible baseline GRAB_DA_ fluorescence prior to stimulation. Each field was imaged once, and 3 to 7 images were captured from each slice. The duration between stimuli was ≥ 5 min. For analyses, a maximum intensity projection image was generated and the pixels with intensities in the top 75% in the intensity histogram of the image stack were used to generate a region of interest (ROI) in ImageJ/Fiji. GRAB_DA3m_ intensity was measured as the average intensity in the ROI. F_0_ was defined as the average fluorescence over 2 s (20 frames) preceding stimulation (F_0_ in striatum, Munc13 control: 126.3 ± 9.7 arbitrary fluorescence units, 6 fields of view/4 slices/4 mice, Munc13 cKO^DA^: 111.4 ± 0.3, 7/3/3, p = 0.18, as assessed by Mann-Whitney rank sum test; F_0_ in midbrain, Munc13 control: 114.3 ± 0.6, 26/6/3, Munc13 cKO^DA^: 124.9 ± 2.2, 14/6/4, p < 0.01, as assessed by a Mann-Whitney rank sum test) and used to determine ΔF/F_0_. The peak fluorescence value was extracted as the maximum change in ΔF/F_0_ over the same ROI from a single image frame from the time series. Sample heatmaps show the peak ΔF/F_0_ image frame with brightness and contrast adjustments identical across genotypes and brain regions; the display color range was set to illustrate the full range of ΔF/F_0_ signal and the underlying raw ΔF/F_0_ was not saturated.

### Experimental design and statistical analyses

Summary data are presented as mean ± SEM with individual values shown as circles. Statistical significance is denoted as *p < 0.05, **p < 0.01, and ***p < 0.001 in each figure. Statistical testing was done in GraphPad Prism 10. The number of observations is described in each figure legend. For comparison of two groups, datasets were tested for normality with a Shapiro-Wilk test. Normally distributed data were assessed with unpaired Student’s t-tests, and non-normally distributed data with Mann-Whitney rank sum tests. For comparisons of multiple groups and variables, two-way ANOVA was used with individual comparisons made using the post-hoc test listed in the corresponding figure legend. Experiments with genotype comparisons were performed by an experimenter blind to genotype during data acquisition and analyses.

## Notes

### Competing Interest Statement

The authors have declared no competing interest.

